# Management of altered metabolic activity in *Drosophila* model of Huntington’s disease by curcumin

**DOI:** 10.1101/2021.01.29.428750

**Authors:** Kumari Aditi, Akanksha Singh, Mallikarjun N Shakarad, Namita Agrawal

## Abstract

Huntington’s disease (HD) is a devastating polyglutamine (polyQ) disorder characterized by extensive neurodegeneration and metabolic abnormalities at systemic, cellular and intracellular levels. Metabolic alterations in HD manifest as abnormal body weight, dysregulated biomolecule levels, impaired adipocyte functions and defective energy state which exacerbate disease progression and pose acute threat to the health of challenged individuals in form of insulin resistance, cardiovascular disease and energy crisis. To colossally mitigate disease symptoms, we tested the efficacy of curcumin in *Drosophila* model of HD. Curcumin is bioactive component of turmeric (*Curcuma longa* Linn), well-known for its ability to modulate metabolic activities. We found that curcumin effectively managed abnormal body weight, dysregulated lipid content and carbohydrate level in HD flies. In addition, curcumin administration lowered elevated reactive-oxygen-species (ROS) levels in adult adipose tissue of diseased flies, and improved survival and locomotor function in HD flies at advanced disease stage. Altogether, these findings clearly suggest that curcumin efficiently attenuates metabolic derangements in HD flies and can prove beneficial in alleviating the complexities associated with HD. Phytochemicals like curcumin that can regulate multiple targets in complex diseases like HD, with least side-effects and maximum benefits, provide a better hope for the treatment of terminally-ill HD patients.

## 1 INTRODUCTION

HD is a dominantly inherited, late-onset and progressive neurodegenerative disorder characterized by locomotor disturbances, cognitive decline and psychiatric abnormalities. In humans, the causative gene for HD is located on chromosome 4p16.3 which comprises polymorphic trinucleotide cytosine-adenine-guanine (CAG)_n_ repeats coding for ~348 kD huntingtin (Htt) protein. Abnormal expansion of (CAG)_n_ repeats beyond normal range of 35 results in HD in humans (HDCRG, 1993).

HD encompass an array of metabolic abnormalities which include cachexia (Sanberg et al., 1981), persistent energy deficit (Gines et al., 2003), hypothalamic-endocrine disturbances with peripheral hormonal dysregulation (Björkqvist et al., 2006; Popovic et al., 2004), and altered biomolecule metabolism *e.g*. fatty acids, protein, cholesterol, glucose *etc*. (Chaves et al., 2017; Mochel et al., 2007; Underwood et al., 2006; Valenza et al., 2005). Multiple defects are found in peripheral tissue such as skeletal (Chaturvedi et al., 2009), pancreas (Andreassen et al., 2002), adipose (Fain et al., 2001) and cardiac cells (Mihm et al., 2007). The multi-faceted metabolic alterations in conjunction with neurodegeneration aggravate the disease condition and augment risk of developing acute complications such as cardiovascular diseases, diabetes, pneumonia *etc*. in HD patients thereby impacting their life expectancy.

The currently available target-specific synthetic drugs for HD offer marginal relief by suppressing the neurological symptoms but cause numerous side effects (Monroy et al., 2013). In this scenario, testing natural compounds with known therapeutic potential and minimum side-effects appear promising. Keeping this aspect as a major consideration, we tested phytochemical ‘curcumin’, a small polyphenolic compound and bioactive component of turmeric (*Curcuma longa* Linn) with strong antioxidant (Hatcher et al., 2008; Kelsey et al., 2010), anti-inflammatory (Aggarwal and Sung, 2009; Ammon and Wahl, 1991; Hatcher et al., 2008), neuroprotective (Darvesh et al., 2012) and metabolic properties (Ammon and Wahl, 1991; Dixit et al., 1988; Rao et al., 1970) in *Drosophila* model of HD.

Several studies have showcased the efficacy of curcumin in animal models of neurodegeneration such as Alzheimer’s (Frautschy et al., 2001; Yang et al., 2005), Parkinson’s (Siddique et al., 2014) and HD (Anjalika & Agrawal, 2016; Elifani et al., 2019; Hickey et al., 2012). Curcumin is shown to suppress inflammation (Lim et al., 2001), increase lifespan (Siddique et al., 2014), ameliorate behavioural dysfunctions (Anjalika & Agrawal, 2016) and improve disease phenotype (Elifani et al., 2019) in these disease models. Besides alleviating neurodegeneration, curcumin also protects organs from toxicities (Singh et al., 2012) and effectively attunes several incapacitating metabolic conditions such as diabetes (Su et al., 2017) and obesity (Ding et al., 2016; Weisberg et al., 2016).

With pleiotropic effects of curcumin in mitigating diverse diseases, we aimed to assess the efficacy of curcumin in attenuation of metabolic derangements linked with HD progression. *Drosophila* model of HD expressing exon1 fragment of human huntingtin with expanded 93 glutamine repeats (Httex1p Q93) in all the neuronal populations recapitulates characteristic features of disease including late-onset nature, mutant Htt accumulation, progressive locomotor dysfunction, photoreceptor neurons degeneration and reduced lifespan (Marsh et al., 2003; Marsh & Thompson, 2004). The diseased flies also exhibit significant alterations in their body weight with extensive modulation in major energy reserves with HD progression (Aditi et al., 2016). Therefore, in order to assuage these symptoms involved in HD pathogenesis, we tested specific doses of curcumin in these flies and evaluated the phytochemical’s efficacy to suppress behavioural and metabolic abnormalities in HD flies.

By dosage studies, we found that an effective concentration of curcumin significantly attenuated dysregulated body weight and total lipid content in diseased flies at specific ages. Additionally, quantification of carbohydrate levels suggested that the same dose effectively regulated circulating trehalose levels and lowered elevated ROS level in adult adipose tissue of diseased flies. Interestingly, the feeding behaviour of diseased larvae or flies reared on curcumin-supplemented food remain unchanged and matched those raised on normal food. Curcumin intake also resulted in an improvement in the survival and locomotor function (data not shown) of diseased flies. Furthermore, expression profiling of metabolic regulators such as dSREBP or HLH106, *bmm* and *lipin* genes revealed no change in their levels in diseased flies upon curcumin supplementation. Altogether, these findings suggest that curcumin proves immensely beneficial in the management of metabolic alterations and energetics in *Drosophila* model of HD. We, therefore, propose that curcumin can serve to be a safe and effective putative therapeutic option for the treatment of HD and will be a boon in the amelioration of HD complexities with minimum side effects. Although, more emphasis will be required to elucidate the complex mechanism of curcumin actions.

## 2 MATERIALS AND METHODS

### 2.1 *Drosophila* stocks, crosses and dietary condition

Transgenic fly lines *w*; P{UAS-Httex1p Q20}, *w*; P{UAS*-*Httex1p Q93}4F1 (Steffan et al., 2001) and *w*; P{w^+mW^.^hs^=GawB}*elav*C155 were used in this study. Expression of polyQ containing peptides was carried out using the bipartite UAS-GAL4 expression system (Marsh et al., 2000). Virgins from stocks with polyQ constructs under the control of yeast UAS were crossed with males from driver line expressing yeast GAL4 transcriptional activator under neuronal *elav* promotor. The resulting progeny expressing polyQ peptides were reared on food mixed without (control) and with 10, 15 and 20µM concentration of curcumin (Sigma Aldrich), with equal supplementation of dimethyl sulfoxide (DMSO, Sigma Aldrich) in all the conditions. Cultures were grown at 25 °C and 65% humidity under a 12 h:12 h light/dark cycle.

### 2.2 Survival assay

For survival analysis, a cohort of freshly emerged flies from control and experimental conditions were collected and transferred to food without and with 10μM concentration of curcumin. Flies were transferred to fresh vials every alternate day till 15 days and mortality in each condition was scored. Minimum 100 flies from each condition were used for the assay. Survival was analysed using Kaplan-Meier estimator.

### 2.3 Food intake assay

The rate of food intake at larval and adult stages was ascertained using colorimetric dye intake assay. Synchronized feeding third instar female larvae were sorted and placed in a hole created at the centre of an agar plate (3.3% wt/vol) filled with yeast paste containing 4% (w/v) FD&C Blue dye # 1 (Sigma Aldrich). The larvae were allowed to feed in the dyed-yeast paste for 2 hours. After feeding, larvae were thoroughly washed with ice cold distilled water, pat-dried and homogenized in 200μl of 1X PBS. The homogenate was centrifuged at 13,500 rpm for 10 minutes and the absorbance of supernatant was recorded at 625 nm. For each condition, two replicates of 10 larvae were used for the assay.

For food intake estimation at adult stages, age matched flies from different experimental conditions were collected and transferred to food containing 2.5% (w/v) FD&C Blue dye # 1 for 30 minutes of feeding. After feeding, the crop and midgut region was dissected and homogenized in 200μl of ice-cold 1X PBS. The homogenate was centrifuged at 13,500 rpm for 10 minutes and absorbance of supernatant was recorded at 625 nm. Three replicates of 10 flies per condition were used and food intake was assessed at day 6, 8 and 12.

### 2.4 Protein quantification

Total protein content was estimated using Bicinchoninic acid (BCA) method. 4 flies/replicate were homogenized in 400 μl of 2% Na_2_SO_4_ and 0.05% Tween 20 (1:1). 80 μl of homogenate was aliquoted in fresh tubes and 500 μl of 0.15% sodium deoxycholate was added. The mixture was left on ice for 10 minutes and 1 ml of 3M trichloroacetic acid was added. The samples were spun at 8,500 rpm for 15 minutes at 4 °C. After discarding the supernatant, pellet was rinsed once with 1 ml of 1M HCL, dried and mixed with 1.6 ml of BCA (Sigma-Aldrich) standard working reagent. The mixture was then incubated at 60 °C for 10 minutes and cooled on ice to stop further colour development. 100 μl of the supernatant was aliquoted and absorbance was recorded at 562 nm. BSA (1μg/μl) was used as standard to calculate protein concentrations. For each condition, three replicates were used for protein estimation.

### 2.5 Glycogen estimation

For glycogen quantification (Xu et al., 2012), 4 flies/replicate were homogenized in 400 μl of 2% Na_2_SO_4_. 20 μl of homogenate was aliquoted and mixed with 46 μl of 2% Na_2_SO_4_ and 934 μl of chloroform/methanol (1:1). The tubes were spun at 13,500 rpm for 10 minutes to obtain glycogen containing pellets which was air dried for 10 minutes. 500 μl of 0.2% (w/v) anthrone reagent (0.2% anthrone in 72% sulphuric acid) (Sigma Aldrich) was added and the mixture was heated at 90 °C for 20 minutes with mixing every 5 minutes. Samples were then cooled on ice for 10 minutes and returned to room temperature for 20 minutes. 100 μl of the supernatant was aliquoted and absorbance was recorded at 620 nm. D-glucose was used as standard to calculate carbohydrate concentration. Five replicates per condition were used for glycogen estimation.

### 2.6 Trehalose estimation

To measure trehalose content (Xu et al., 2012), 4 flies/replicate were homogenized in 500 μl of 70% ethanol and tubes were spun at 5,000 rpm at 4 °C. The pellets were resuspended in 200 μl of 2% NaOH, heated at 100 °C for 10 minutes and cooled on ice. 100 μl of mixture was then aliquoted and 750 μl of anthrone reagent was added. The samples were heated at 90 °C for 20 minutes with mixing every 5 minutes, cooled on ice for 10 minutes and returned to room temperature for 20 minutes. 100 μl of the supernatant was aliquoted and absorbance was recorded at 620 nm. Trehalose concentration was calculated using D-glucose standard. For each condition, five replicates were used for the assay.

### 2.7 Lipid estimation

For each condition, groups of 10 freshly emerged flies were collected, etherized and weighed immediately in micrograms using Citizen CM11 microbalance to obtain fresh weight. The samples were dried afterwards in a preheated oven at 70 °C for 36 hours and weighed again to obtain dry weight. The difference between fresh weight and dry weight represented water content of the flies. Subsequently, ether soluble lipids were extracted from the samples by adding 1 ml of diethyl ether to intact dry flies stored in 1.5 ml microcentrifuge tubes. The tubes were kept on a gel rocker and lipid extraction was carried out for 48 hours with three ether changes at an interval of 12 hours each. After the last ether change, flies were dried for 2 hours at 30-35 °C and weighed again to obtain lipid free weight of flies. Lipid content of the flies was calculated by subtracting lipid free weight from dry weight (Chandrashekara et al., 2014; Handa et al., 2014). Five replicates per condition were used and lipid content was assessed at 0, 3, 5, 7, 9, 11 and 13 days of age.

### 2.8 Lipid droplet staining

For lipid droplet staining in adult adipose tissue, age-matched flies were collected, etherized and their abdomen was dissected in ice-cold 1X PBS. The abdomens were fixed in 4% formaldehyde in PBT for 20 minutes at room temperature. After fixation, sheets of fat body were detached from the dorsal abdominal area, fixed for additional 10 minutes and washed thrice for 10 minutes each with 1X PBS. The tissue was stained with 1:2000 dilution of 0.5 mg/ml Nile Red (Sigma Aldrich, N3013) in PBS for 30 minutes. After staining, the tissue was rinsed twice with 1X PBS and mounted in Vectashield mounting medium (Vector Labs, H-1000). Images were acquired using Nikon Eclipse (Ni-E) fluorescence microscope and lipid droplets quantification was performed using NIS-Elements AR software. Minimum five to six samples per condition were analyzed for lipid droplets quantification.

### 2.9 Reactive oxygen species (ROS) detection and imaging

Dihydroethidium (DHE) staining was used to monitor superoxide radicals in the adult adipose tissue of flies from different experimental conditions (Owusu-Ansah et al., 2008). Age-matched flies were etherized and adult fat body was dissected from the dorsal surface of abdomen. To allow optimal respiration, tissue was dissected in 1X Schneider’s *Drosophila* medium (+) L-glutamine (GIBCO, 21720024) and any extraneous tissue was removed. A stock solution of 30 mM DHE (Invitrogen Molecular Probes, D11347) was reconstituted just before use in anhydrous DMSO (Sigma-Aldrich, 276855). For staining, the reconstituted dye was further diluted in 1X Schneider’s medium to obtain a final working concentration of 50 μM and vortexed to disperse the dye evenly. The tissue was incubated with the dye for 4 minutes in dark soon after dissection at room temperature. The samples were rinsed twice in 1X Schneider’s medium and images were acquired immediately using Nikon Eclipse (Ni-E) fluorescence microscope. ROS quantification was performed using NIS-Elements AR software. Minimum six to seven samples per condition were analysed for ROS quantification.

### 2.10 Quantitative Real Time PCR expression profiling

For transcriptional analysis, flies from different experimental conditions were collected, snap frozen and stored at −80 °C until processing. Total RNA from these flies was isolated using TRIZOL® Reagent (Thermo Fisher Scientific, 15596-026) following standard protocol of RNA extraction from *Drosophila* (Auburn, 2006). After RNA extraction, 1 μg of the total RNA was used for cDNA synthesis with Revert Aid First Strand cDNA Synthesis Kit (Thermo Fisher Scientific, K1622). Thereafter, qRT-PCR reaction was performed on CFX Connect Real-Time System (Bio-Rad Laboratories) using SYBR-Green based detection system with QuantiNova SYBR green PCR Kit (Qiagen, 208052). Relative expression of mRNA was calculated through ΔΔCt method using the ribosomal protein RP49 RNA expression as internal control for each sample. Primers used in the qRT-PCR analysis were designed online using Primer3 software and evaluated on NCBI-Blast. Minimum 6 replicates per condition with 6 flies per replicate were used for the assay.

### 2.11 Statistical analysis

Statistical analysis was performed in IBM SPSS statistical package (version 22). The results are represented as mean ± S.E.M in all the graphs. Normality and homoscedasticity of the data was assessed using Shapiro-Wilk and Levene’s test, respectively. Data was analysed using Student’s t-test and multi-factor analysis of variance (ANOVA) followed by Tukey HSD post-hoc test for multiple comparisons and Test of Simple Effects for pairwise comparisons. Survival data was analysed using Kaplan-Meier test.

## 3 RESULTS

### 3.1 Curcumin supplementation improves median survival in HD flies

In the present study, a dosage effect of curcumin *i.e*. 0, 10, 15 and 20μM was assessed on the locomotor performance as well as survival of HD *Drosophila*. A cohort of female flies expressing unexpanded (Httex1p Q20) and expanded glutamine repeats (Httex1p Q93) was collected and reared on food supplemented without and with different concentration of curcumin. Initially, their vertical locomotor performance was assessed at day 1, 3, 7 and 9 post-eclosion. Significant amelioration in behavioural dysfunction with curcumin supplementation in 7-day old HD flies has previously been reported (Anjalika & Agrawal, 2016), however in this study, we chose an additional time-point *i.e*. day 9 to see whether curcumin shows beneficial effect at later stage when both disease symptoms and metabolic dysregulation become more distressing. We found that among different tested concentrations, 10μM curcumin fed diseased flies display significant improvement in their motor activity at day 7 and 9 (Tukey HSDα0.05, day 7, n = 20; p = 0.000073; day 9, n = 20; p = 0.023) as compared to the age-matched diseased flies reared on normal food (data not shown).

We further determined survivorship of diseased flies in response to 10μM curcumin treatment. Lifespan and survival are multifaceted processes which can be influenced by several factors such as metabolic state, diet, physical activity, underlying disease etc. After survival analysis, we found that although the survival curve of unfed (Httex1p Q93) and curcumin-fed diseased flies remain comparable to each other [log-rank test, Httex1p Q93 *vs* Httex1p Q93 (10μM curcumin), *p* = 0.3049], the median survival of diseased flies receiving 10μM dose of curcumin improved from ~5 days (unfed) to ~7 days. Also, ~75% death time of these flies was found to be increased *i.e.* ~13 days as compared to that of 11 days in unfed flies. Therefore, at advanced stage of HD, 10μM dose of curcumin proved quite effective in improving the survival of diseased flies as compared to those receiving normal food. The control flies (Httex1p Q20) exhibited no signs of mortality up to 15 days, suggesting that the transgene with unexpanded glutamine had no notable toxic effect (Figure 1). Altogether, these results suggest that 10μM dose of curcumin efficiently improves survival as well as locomotor dysfunction in diseased flies even at terminal stage when the disease results in both behavioural and physiological deterioration.

**Figure 1.**
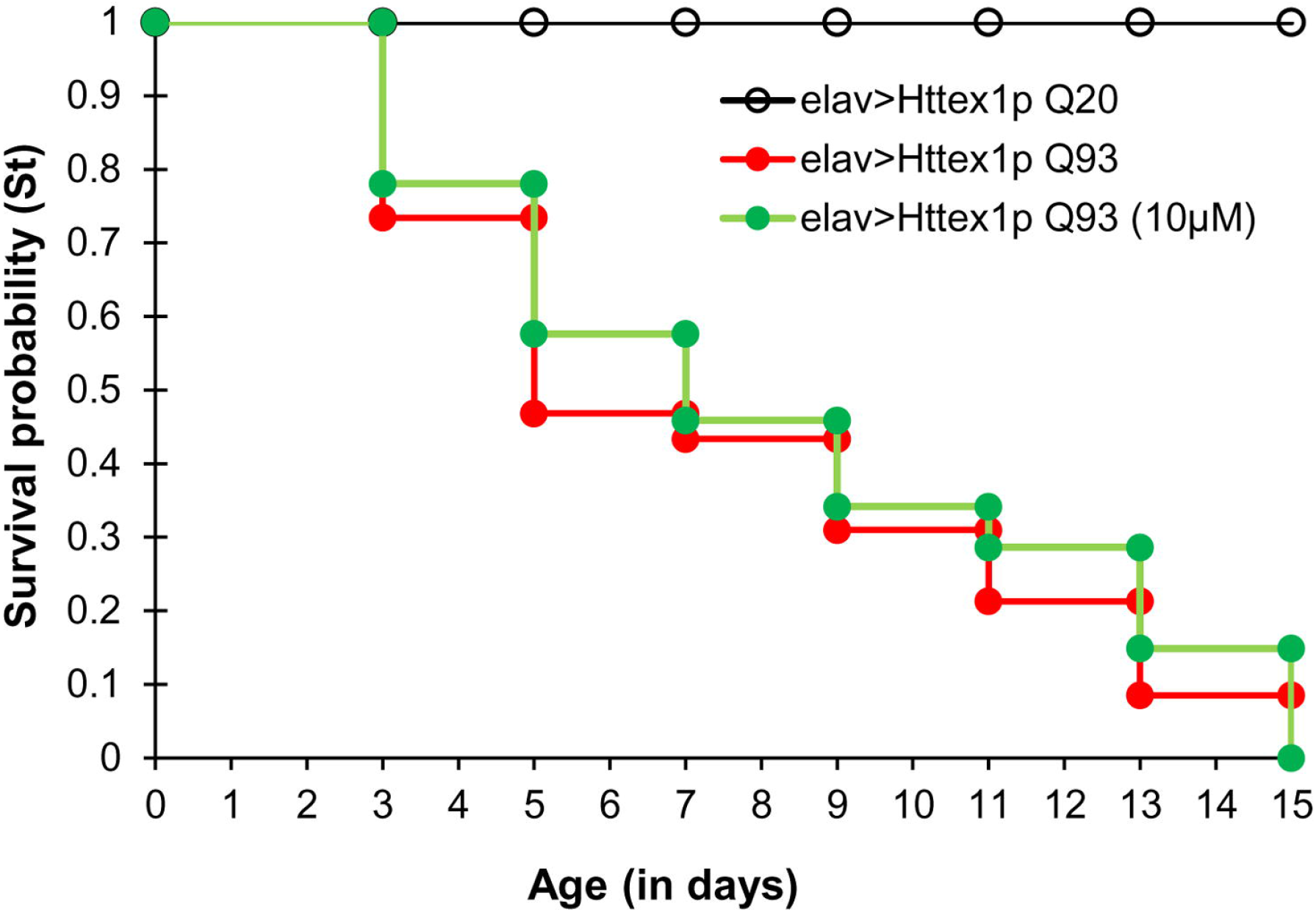
Curcumin improves survival in HD flies. Survival probability of flies with unexpanded (elav>Httex1p Q20) and expanded glutamine repeats (elav>Httex1pQ93) fed without and with 10μM concentration of curcumin from larval stages. Notable improvement was observed in the median survival time of 10μM curcumin fed diseased flies, as compared to the unfed controls. Median survival (50%) is ~5 days for elav>Httex1pQ93 which increased upto ~7 days for 10μM curcumin fed diseased flies. Interestingly, 75% death time for diseased condition is just ~11 days whereas it extends upto ~13 days for 10μM curcumin fed diseased flies. Survival data was analysed using Kaplan-Meier method followed by log-rank test (elav>Httex1p Q93 vs. elav>Httex1p Q93 (10μM), *p* = 0.3049). For each condition, n = 100.

### 3.2 Curcumin manages altered body weight, dry mass and water content in HD flies

Previously, we have reported extensive metabolic alterations in HD flies as a consequence of neuronal expression of mutant Htt (Aditi et al., 2016). Regulation of metabolism can possibly muffle HD symptoms, particularly at advanced stages of the disease. HD flies exhibit significant weight modulation during entire course of the disease with excessive increase in weight during disease onset followed by significant reduction at terminal stage (Aditi et al., 2016). In an attempt to test the effect of curcumin in body weight management, newly-emerged normal and diseased flies were reared on food supplemented without and with 10μM dose of curcumin from early larval stage till 13 days post-eclosion. Quantification of fresh body weight showed that 10μM dose of curcumin significantly attenuates body weight alterations observed in HD flies not fed with curcumin with disease progression. Diseased flies fed with 10μM curcumin exhibited a significant decline in their abnormally high body weight at day 3 (Tukey HSD_α0.05_, *n* = 50, *p* = 0.001) and 7 (Tukey HSD_α0.05_, *n* = 50, *p* = 0.000041) followed by significant improvement in deteriorating body weight at day 11 (Tukey HSD_α0.05_, *n* = 50, *p* = 0.000209) as compared to the age-matched diseased flies. Interestingly, we observed that body weight of diseased flies reared on 10μM curcumin became comparable to age-matched flies with unexpanded glutamines at day 3, 7 and 11. Flies with unexpanded glutamines did not display any variation in their body weight upon receiving the same dose of curcumin for 13 days (Figure 2a). Body weight of an organism is majorly constituted by water, protein, carbohydrates and lipids, hence, further understanding of the relative proportions of these molecules in control and curcumin-fed HD flies will help gain a deeper insight.

**Figure 2.**
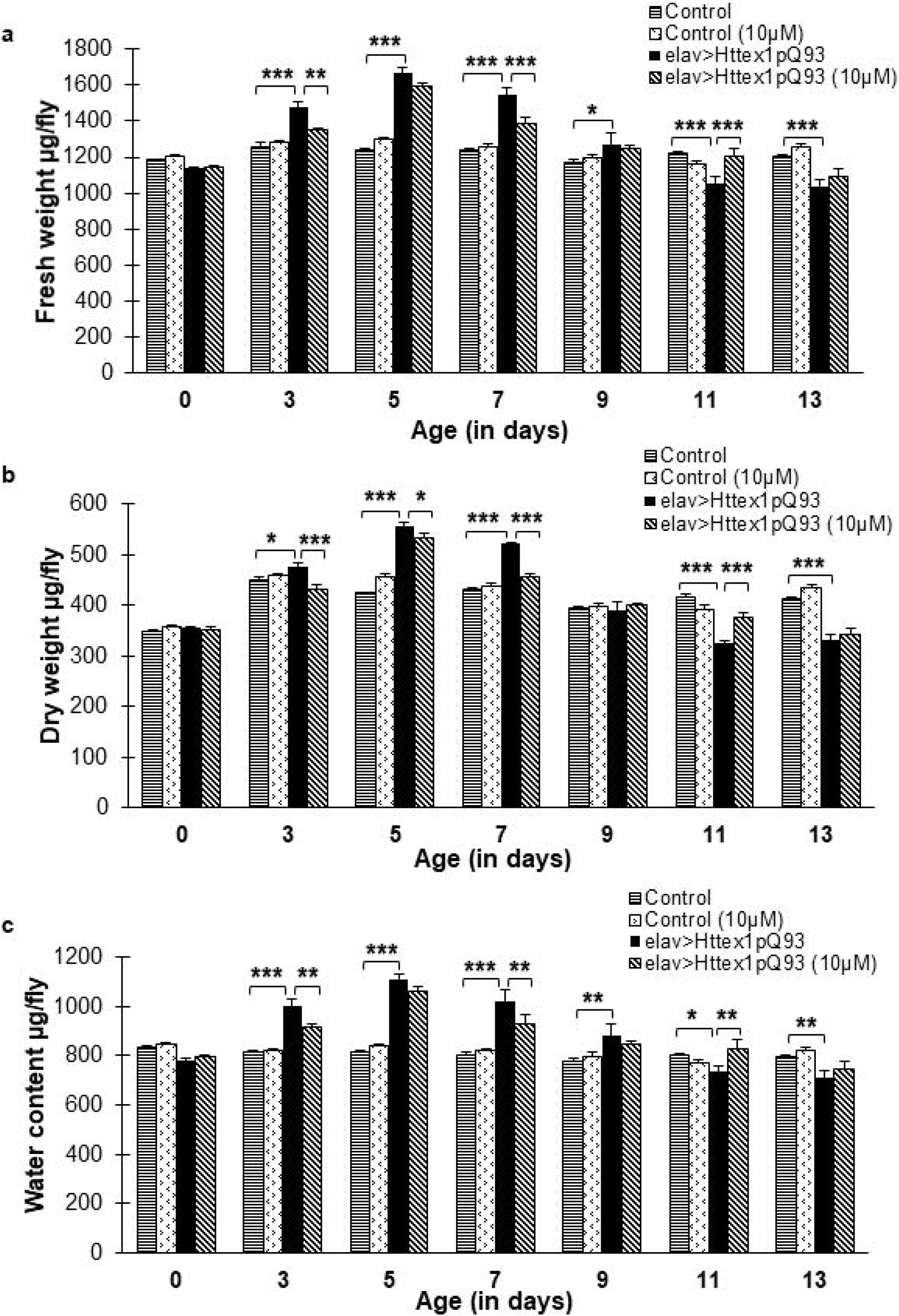
Dietary curcumin modulates body weight, dry mass and water content. Curcumin administration significantly regulates abnormally high and low body weight (a) at day 3, 7, and 11 (b) dry mass at day 3, 5 and 7 in diseased flies (c) dysregulated water content is managed at day 3, 7 and 13 in diseased flies upon curcumin feeding. Data was analyzed using multi-factor ANOVA followed by Tukey HSD post hoc test and Test of Simple Effects. Values are represented as mean ± S.E.M. Tukey HSD_α0.05_, *** *p <* 0.001; ** *p <* 0.01; * *p* < 0.05. For each condition, *n* = 50.

We assessed the impact of curcumin on dry weight and water content of HD flies. Dry weight assessment revealed that administration of 10μM dose of curcumin significantly amend abnormally high dry weight in disease flies at day 3 (Tukey HSD_α0.05_, *n* = 50, *p* = 0.000032), 5 (Tukey HSD_α0.05_, *n* = 50, *p* = 0.030) and 7 (Tukey HSD_α0.05_, *n* = 50, *p* = 2.6012E-9), followed by significant improvement in severely low dry mass at day 11 (Tukey HSD_α0.05_, *n* = 50, *p* = 3.00E-6), as compared to the untreated diseased flies. Previously, we had reported significant alterations in dry weight of HD flies through the entire course of disease (Aditi et al., 2016). Flies with unexpanded glutamines receiving 10μM curcumin however exhibited modulation in dry weight at day 5 (Tukey HSD_α0.05_, *n* = 50, *p* = 0.002), 11 (Tukey HSD_α0.05_, *n* = 50, *p* = 0.018) and 13 (Tukey HSD_α0.05_, *n* = 50, *p* = 0.028) as compared to the age-matched control flies (Figure 2b).

Evaluation of water content revealed that diseased flies exhibited an altered pattern of water level from day 3 to 13 (Tukey HSD_α0.05_, *n* = 50; day 3, *p* = 1.634E-8; day 5, *p* = 3.567E-16; day 7, *p* = 1.223E-10, day 9, *p* = 0.001182; day 11, *p* = 0.03236; day 13, *p* = 0.00494), when compared to age-matched unexpanded flies, as reported previously (Aditi et al., 2016). Interestingly, when fed with 10μM dose of curcumin, diseased flies exhibited a significant reduction in their elevated water levels at day 3 (Tukey HSD_α0.05_, *n* = 50; *p* = 0.009255) and 7 (Tukey HSD_α0.05_, *n* = 50; *p* = 0.0033913), followed by a significant improvement at day 11 (Tukey HSD_α0.05_, *n* = 50; *p* = 0.0033980) the period when their water content decline below normal levels (Figure 2c). No detectable effect of curcumin was observed on the water content of normal flies with unexpanded glutamine from day 0 till day 13. Modulation of dry weight and water content of HD flies by curcumin suggests that it is perhaps micromanaging the metabolic and/or catabolic processes in diseased condition.

### 3.3 Dietary curcumin has no effect on food intake in diseased flies

Feeding is an inherent characteristic of an organism with profound effect on whole body composition, energy stores, metabolic activity and life span. Food intake quantification becomes critical while assessing behaviour, nutrition and drug administration.

In order to ascertain the probable cause of weight modulation, we quantified food intake in normal and diseased condition, at both larval and adult stages, reared on diet without and with 10μM curcumin. The synchronized feeding stage third instar larvae and adult flies, grown on control or curcumin-supplemented medium were fed with blue dye. As shown previously (Anjalika and Agrawal, 2016), we found that normal and diseased larvae fed with 10uM dose of curcumin show feeding behaviour comparable to those fed diet without curcumin (data not shown, two-way ANOVA, *n* = 20, F_1, 4_ = 0.080, *p* = 0.791). Similarly, at adult stage too, there was no difference in the food intake of 6, 8 and 12 day old normal and diseased flies due to curcumin supplementation (Figure 3, multi-factor ANOVA, *n* = 20, F_2, 36_ = 0.110, *p* = 0.896). These results clearly suggest that curcumin modulates body weight in diseased flies either by amendment of systemic metabolism or cellular metabolic components instead of directly altering the feeding response. Therefore, we proceeded to ascertain which of the macromolecules, if any, curcumin modulates by quantifying proteins, glycogen, trehalose and lipid levels in these flies.

**Figure 3.**
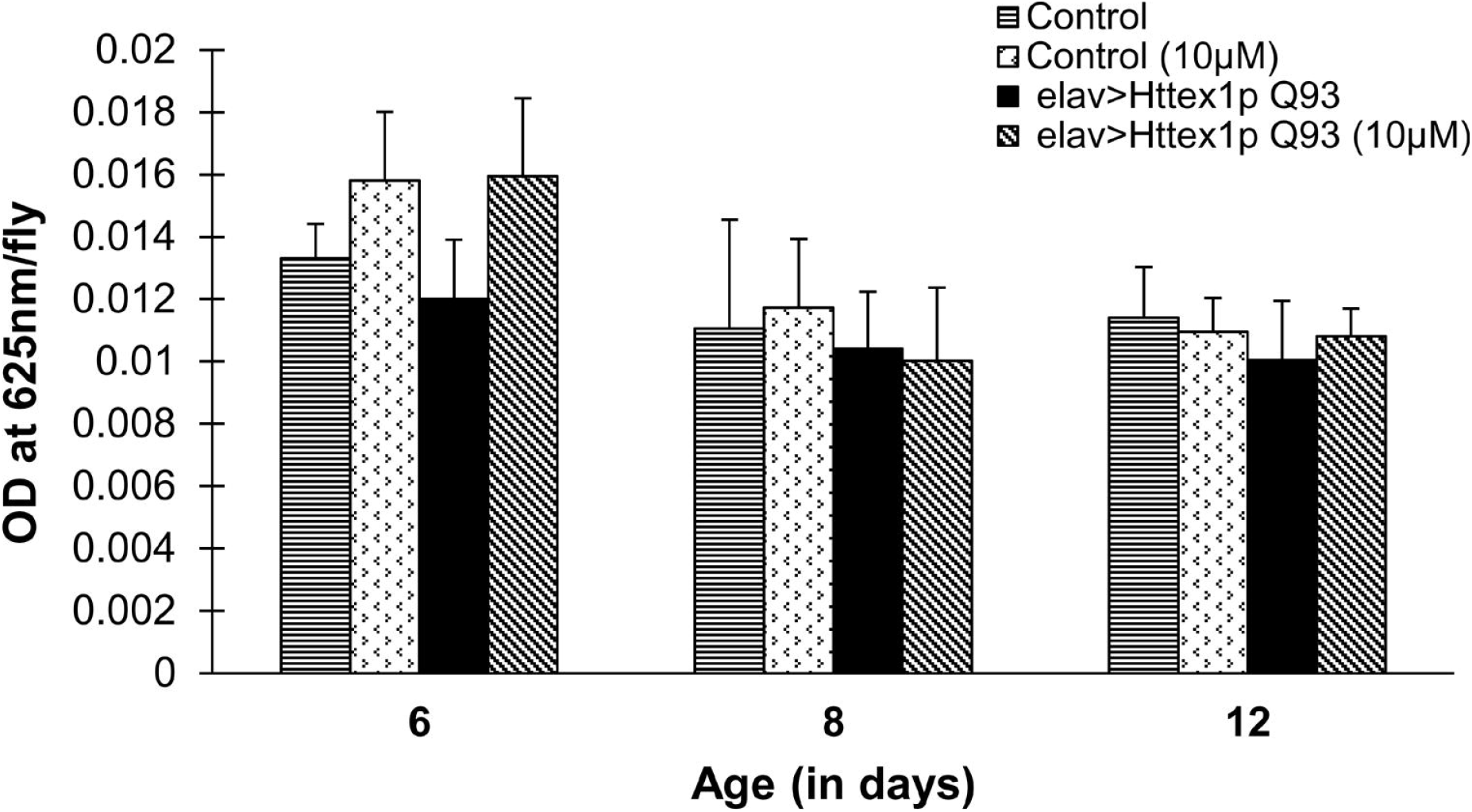
Feeding behavior of HD flies remain unchanged by curcumin supplementation in the food. Feeding of normal (elav>Httex1p Q20) and diseased (elav>Httex1p Q93) female adult flies reared on control or 10μM curcumin supplemented diet was measured using colorimetric dye intake assay. No difference was observed in food intake of 6, 8- and 12-day old control and diseased females supplemented without or with effective concentration of curcumin. Data was analyzed using multi-factor ANOVA, F_2, 36_ = 0.110, *p* = 0.896. Values are represented as mean ± S.E.M and for each group, *n* = 20 flies.

### 3.4 Curcumin superintends macromolecules involved in metabolism

In an attempt to elucidate a detailed account of curcumin action on components involved in metabolism, we investigated the effect of curcumin feeding on total protein, glycogen and circulating disaccharide trehalose levels in both normal and diseased condition without or with curcumin supplementation. Surprisingly, we did not find any effect of 10μM curcumin on the levels of total protein (Figure 4a) and glycogen content (Figure 4b) in normal or diseased flies. However, administration of 10μM dose of curcumin resulted in significant (Tukey HSD_α0.05_, *n* = 20, *p* = 5.00E-6) decrease in the trehalose level in diseased flies at day 7. Further, no notable effect of curcumin on the trehalose levels of age matched flies with unexpanded glutamine, except decrease at day 3 (Tukey HSD_α0.05_, *n* = 50, *p* = 0.034) (Figure 4c) was found. These findings further suggested that curcumin intake has no substantial effect on the protein or stored carbohydrate in unexpanded or HD flies, however, there is a noticeable effect of phytochemical on the circulating sugar trehalose levels at specific ages in both normal and diseased flies.

**Figure 4.**
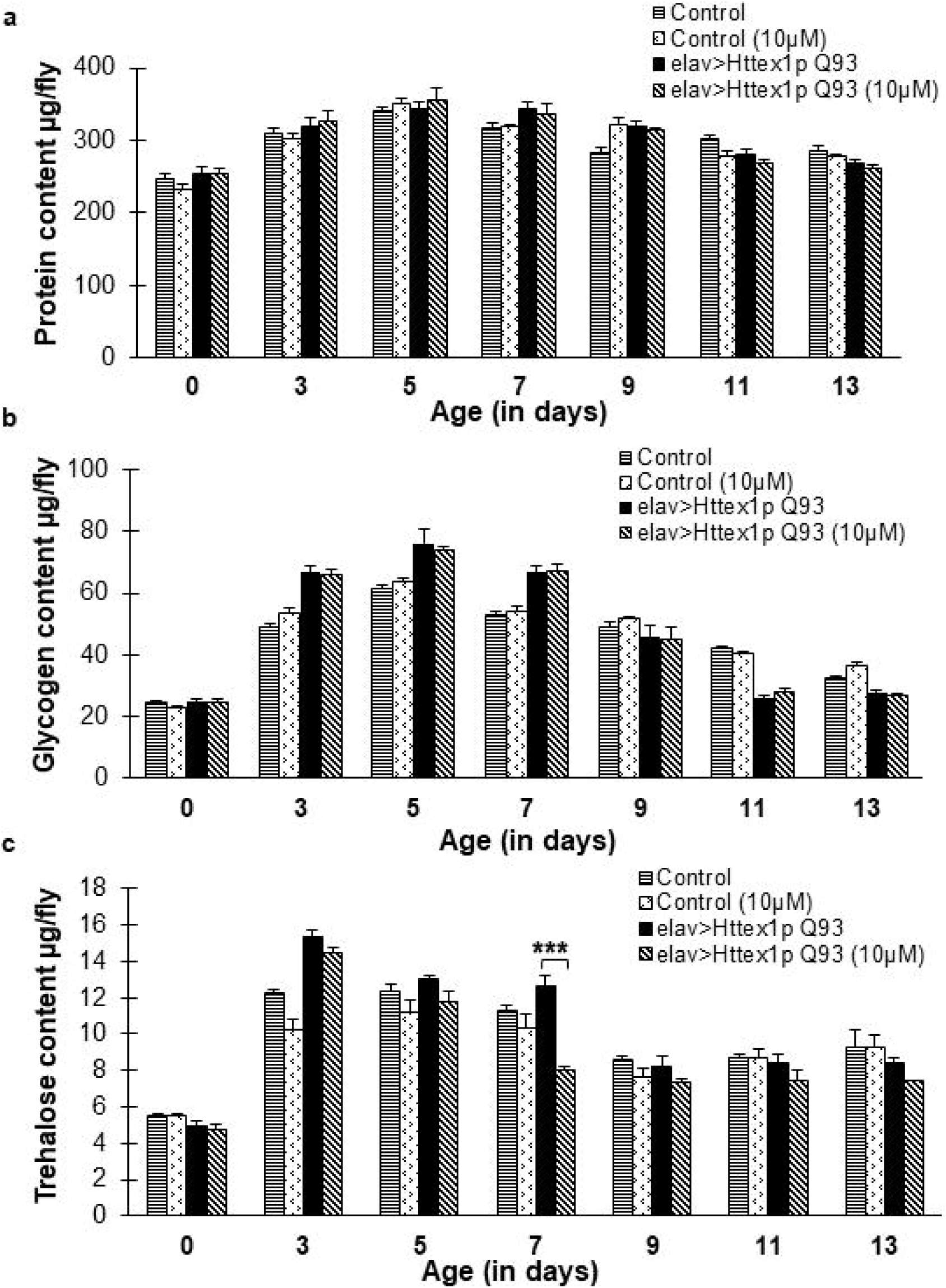
Curcumin regulates circulating sugar trehalose in diseased flies. There is no notable effect of curcumin on the levels of (a) protein, and (b) glycogen in HD flies. On the contrary, (c) significant reduction in otherwise increased levels of circulating sugar trehalose is observed in curcumin fed diseased flies at day 7. Data was analyzed using multi-factor ANOVA followed by Tukey HSD post hoc test and Test of Simple Effects. Values are represented as mean ± S.E.M. Tukey HSD_α0.05_, *** *p <* 0.001; ** *p <* 0.01; * *p* < 0.05. For each condition, *n* = 20 for protein, glycogen and trehalose content quantification.

### 3.5 Curcumin attunes total lipid and subcellular lipid content in HD flies

To unravel the effect of curcumin on major energy reserves, we quantified total lipid content in normal and diseased flies reared on diet without and with 10μM curcumin up to 13 days. Lipids constitute a dominant fraction in organism’s body weight, therefore any modulation in body weight indicates probable regulation in lipid metabolism. We have reported earlier that there is significant perturbation in lipid level in diseased flies through the entire course of disease which becomes initially high and then low (Tukey HSD_α0.05_, *n* = 50; day 5, *p* = 3.942E-11; day 7, *p* = 0.000060; day 9, *p* = 0.00020; day 11, *p* = 0.000080 and day 13, *p* = 0.0000) as compared to age-matched control flies with unexpanded glutamines (Aditi et al., 2016). Interestingly, in the present study with 10μM curcumin supplementation, significant attenuation in dysregulated total lipid content of diseased flies was observed at day 3 (Tukey HSD_α0.05_, *n* = 50, *p* = 0.01682) and upon progression at day 7 (Tukey HSD_α0.05_, *n* = 50, *p* = 4.411E-7). However, decline in lipid levels at advanced disease stage i.e. day 9 (Tukey HSD_α0.05_, *n* = 50, *p* = 0.03493) could not be reinstated by curcumin. We found no notable difference in lipid content by supplementation of curcumin in 0 to 13-day old normal flies with exception at day 5 (Tukey HSD_α0.05_, *n* = 50, *p* = 0.0404; Figure 5a). These results suggest that the administered dose of curcumin modulates altered lipid level in diseased flies at different ages which might be linked to their improved metabolic profile. However, at terminal disease stage as marked by day 11 or 13, the insidious disease progression seems to outweigh the phytochemical’s beneficial effects.

**Figure 5.**
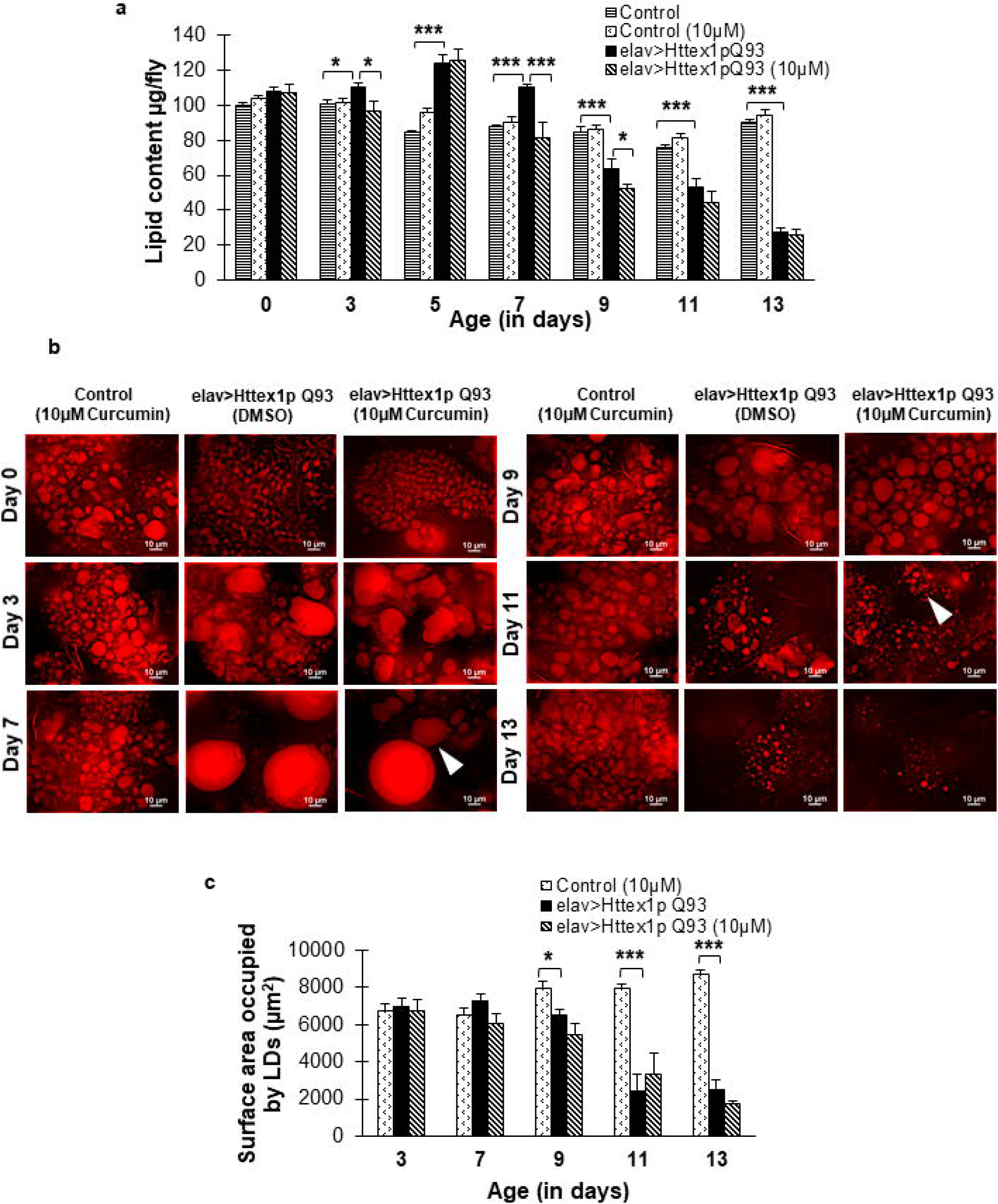
Curcumin modulates total lipid content. (a) Dietary curcumin significantly decreases abnormally high lipid levels in diseased flies at day 3 and 7 which becomes comparable to those of age-matched flies with unexpanded glutamines. However, the lipid levels further decline significantly at day 9 in curcumin fed diseased flies. (b) At sub-cellular level, curcumin intake improves distribution of intracellular lipid in lipid droplets (LDs) in abdominal adipose tissue of diseased flies at day 7 and 11. Arrowheads indicate presence of smaller LDs at day 7 and moderately improved LD distribution at day 11. Scale bar represents 10μm. (c) Quantification of total surface area occupied by LDs suggests a considerable reduction in bigger LDs at day 7 and subsequent improvement in distribution at day 11 by curcumin intake. Lipid content data was analyzed using multi-factor ANOVA followed by Tukey HSD post hoc test and Test of Simple Effects and LD quantification was analyzed using two-way ANOVA. Values are represented as mean ± S.E.M. Tukey HSD_α0.05_, *** *p <* 0.001; ** *p <* 0.01; * *p <* 0.05. For each condition, *n* = 50 for total lipid content and *n* = 6 for LD quantification.

Besides estimating total lipid content, to ascertain the probable effect of curcumin on subcellular lipids, we monitored distribution of lipid droplets (LDs), the storehouse of intracellular lipids which enrich *Drosophila* adipocytes. LDs are highly dynamic organelles with hydrophobic core comprising of neutral triacylglycerols (TAGs). We found that there was a noticeable regulation in LDs distribution in disease condition after supplementation of 10μM curcumin. Large LDs present in adipose tissue in 7-day old HD flies exhibited marked reduction in their size with 10μM curcumin administration. In addition, at day 11, LD distribution improved upon curcumin supplementation as compared to untreated flies with expanded glutamines (Figure 5b, c). However, 10μM dose of curcumin had no observable effect on the abundance of LDs in adipose tissue of flies with unexpanded glutamines. Taken together, these results suggest that 10μM dose of curcumin exhibits beneficial effects on the altered intracellular lipid abundance that ultimately reflects in the modulation of total lipid content and may have a beneficial impact on the maintenance of overall metabolic homeostasis in HD flies.

### 3.6 Curcumin suppresses oxidative stress in diseased condition

Curcumin acts as an excellent antioxidant and metabolic regulator, and by our results it is evident that curcumin effectively improves the overall metabolic condition and maintains energy balance, likely by impacting vital organs. *Drosophila* fat body is a key metabolic organ which performs multiple functions such as nutrient storage and mobilization, thus, we investigated the effect of curcumin on the fat body in normal and HD flies and monitored their ROS levels.

Basal ROS level was detected in the adipose tissue of 7- and 13-day old flies with unexpanded glutamines whereas ROS in abdominal fat body of 7 and 13 (Tukey HSD_α0.05_, *n* = 7, *p* = 0.024) day old diseased flies was considerably higher as compared to age-matched control flies. Further, diseased flies displayed significantly high ROS at day 13 (Tukey HSD_α0.05_, *n* = 7, *p* = 0.010) in comparison to 7-day old counterparts which clearly indicated that ROS levels increase with disease progression. Interestingly, significant decline in the elevated ROS levels in adult adipose tissue of diseased flies reared on 10μM curcumin was found at day 13 (Tukey HSD_α0.05_, *n* = 7, *p* = 0.021) as compared to those reared without curcumin (Figure 6a, b).

**Figure 6.**
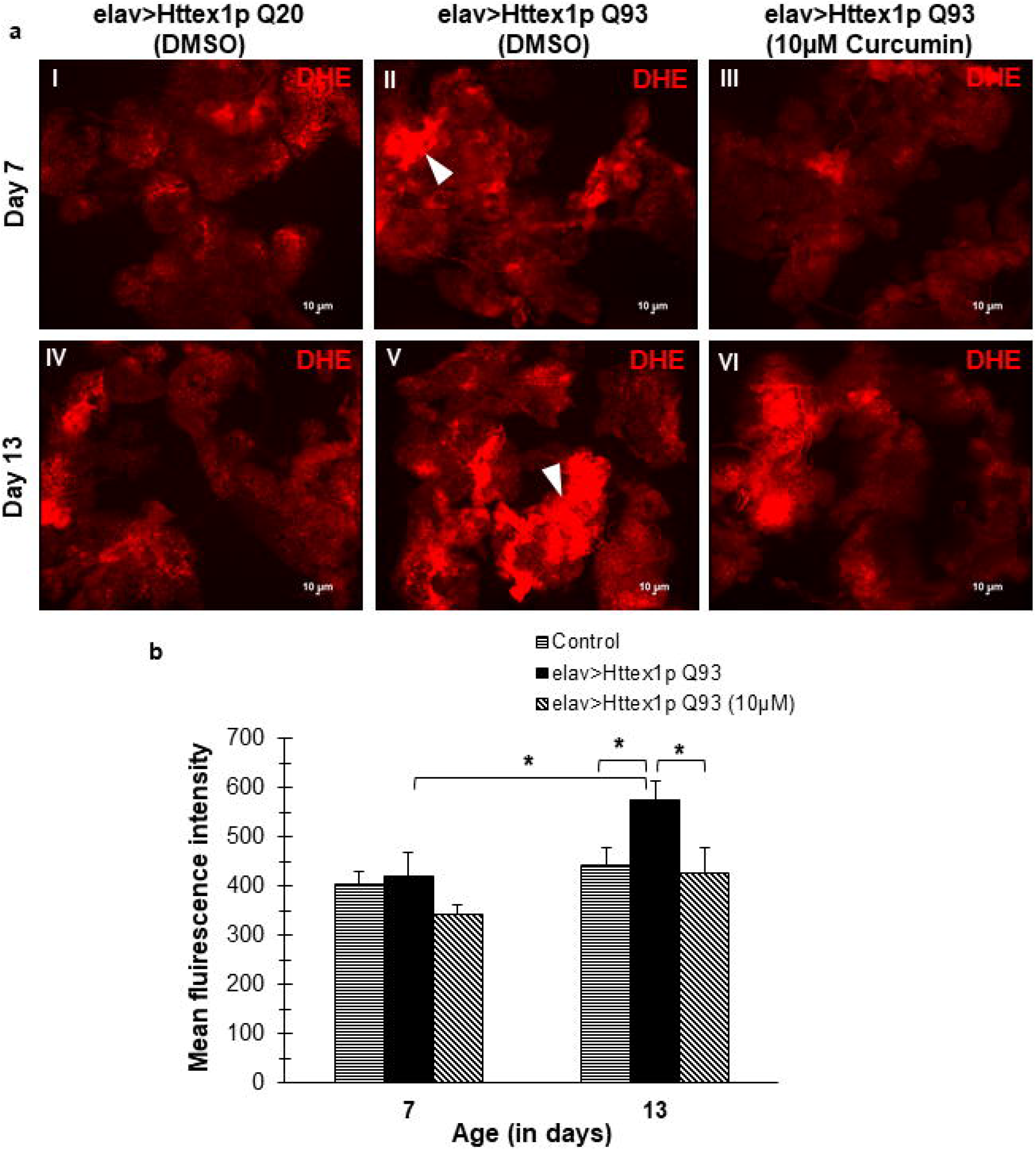
Curcumin reduces elevated ROS levels in adipose tissue of diseased flies. (a, I-VI) Evaluation of ROS production in abdominal fat body of 7 and 13-day old normal females (elav>Httex1p Q20; panels I, IV), diseased (elav>Httex1p Q93; panels II, V), and diseased flies supplemented with 10μM curcumin (panel III, VI). Adult fat body from 7 and 13-day old diseased condition displayed increased levels of ROS as compared to age-matched normal flies. Interestingly, curcumin intake in diseased condition suppressed elevated levels of ROS in diseased condition. Arrowheads show areas in adult fat body producing higher ROS levels than neighbouring cells. Red = dihydroethidium (DHE), scale bars represent 10μm. (B) Quantification of mean fluorescence intensity of DHE staining in adult fat body of 7 and 13 day old normal, diseased and curcumin fed flies revealed significant increase in ROS intensity in day 13 diseased condition as compared to day 7 diseased or age-matched normal flies. Furthermore, there was a significant effect of curcumin treatment on ROS intensity levels (F_1, 24_ = 8.782, *p* = 0.007). Curcumin feeding results in significant decrease in elevated ROS levels in day 13 old diseased flies which becomes comparable to those of age-matched normal flies. Data was analysed using two-way ANOVA followed by Tukey HSD post hoc test. Values are represented as mean ± S.E.M. Tukey HSD_α0.05_, \*\**p* < 0.01. For each condition, *n* = 7 female flies.

Collectively, these results indicated that high ROS levels detected in the adipose tissue in 7 and 13-day old diseased flies might underlie the altered cellular functions that scale up to alter tissue function and in turn contribute to the persistent disruptions in systemic metabolic state as evident in HD flies. Administration of 10μM curcumin lowers the free radical levels in adipose tissue at advanced stage of the disease thereby providing protection against increased oxidative insult. This may ultimately result in the fine tuning of major biomolecules and amelioration of disease symptoms in HD flies.

### 3.7 dSREBP, *bmm* and *lipin* levels in curcumin-fed HD flies match with control

Diseased flies showcased high ROS in adult adipose tissue which might deter the optimal organ functioning. Generally, ROS production and lipid metabolism pathways remain intricately entwined, and an abnormal lipid level may lead to increased oxidative stress and *vice versa*, ultimately disrupting the overall cellular performance. Therefore, improved ROS and lipid levels point towards a probable regulation of key metabolic regulators in curcumin-fed HD flies. dSREBP (HLH106), *bmm* and *lipin* are few key effectors genes which regulate lipogenesis, lipolysis, energy metabolism and inflammation in *Drosophila*.

To decipher the basis of curcumin action in diseased condition, we monitored the expression of these genes in normal and diseased flies reared without and with curcumin. There was no change in the mRNA level of dSREBP, *bmm* and *lipin* genes in diseased flies at day 7 and 13 or with 10μM curcumin administration (Figure 7). The variation in lipid or ROS levels did not correlate with the expression level of dSREBP, *bmm* or *lipin* genes. In normal flies, however, we observed modulation in dSREBP expression with curcumin supplementation (Figure 7a). Significant decrease at day 7 (student’s t-test, *n* = 36, *p* = 0.00454) followed by significant upregulation at day 13 (student’s t-test, *n* = 36, *p* = 0.0195) in dSREBP mRNA levels was seen in curcumin-fed normal flies, but they did not project to their corresponding lipid levels at day 7 or 13.

**Figure 7.**
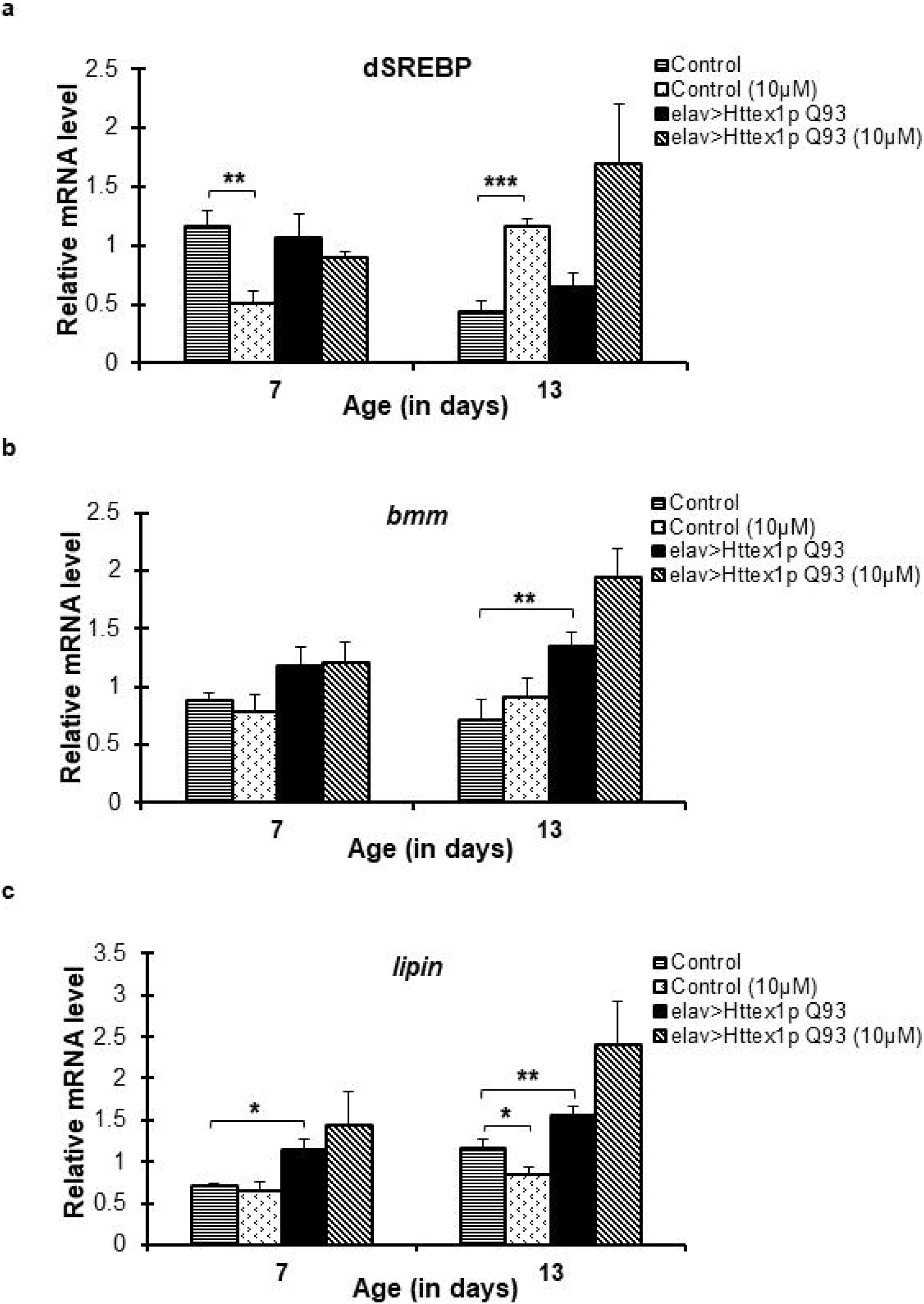
dSREBP, *bmm* and *lipin* expression remain unmodulated in HD flies by curcumin supplementation. (a) dSREBP or HLH106 mRNA levels in 7 and 13-day old normal (elav>Httex1p Q20) and diseased (elav>Httex1p Q93) flies reared on control or 10μM curcumin supplemented diet was monitored using quantitative RT-PCR. Normal flies exhibited significant decrease in dSREBP mRNA levels at day 7 followed by significant increase at day 13, whereas, diseased flies did not display any change in the expression of dSREBP gene at both the ages without or with 10μM curcumin supplementation. (b) *bmm* mRNA levels remain unchanged in 7 and 13 day old normal and diseased flies reared without and with 10μM curcumin. (c) No change in *lipin* mRNA levels was seen in 7 or 13-day old diseased flies reared on 10μM curcumin diet. Normal flies showed significant decrease in *lipin* mRNA at day 13 upon curcumin feeding. Data was analyzed using student’s t-test. Values are represented as mean ± S.E.M. *** *p <* 0.001; ** *p <* 0.01. Sample size: 6 flies/replicate, 6 replicates/condition.

No change in *bmm* mRNA was observed either in normal or diseased flies, except at day 13 when diseased flies exhibited higher *bmm* mRNA levels in comparison to age-matched normal flies (Figure 7b). However, no effect of curcumin on *bmm* expression was seen. Similarly, HD flies fed with curcumin did not show any modulation in their *lipin* expression at any age. Conversely, untreated diseased flies exhibited significantly high *lipin* mRNA levels at both day 7 and 13 as compared to control flies. Normal flies also showed significant decline in *lipin* mRNA levels at day 13 with curcumin supplementation (Figure 7c). These results clearly indicated that curcumin did not modulate the action of these three genes in HD.

## 4 DISCUSSION

Turmeric has a long history of traditional use in Asian population as food additive, herbal remedy and medicine (Ammon and Wahl, 1991; Hatcher et al., 2008). Asian population also have the lowest prevalence of HD at 0.40/100,000, as compared to a high prevalence of 2.71/100,000 worldwide (Pringsheim et al., 2012). The toxicological profile of curcumin deems it extremely safe and it can be used a regular part of diet and medicine (Ganiger et al., 2007). Several *in vitro* and *in vivo* studies have confirmed the activity of curcumin in biological system by either detecting curcumin or its bio-transformed metabolites in plasma, peripheral organs and brain (Begum et al., 2008; Ireson et al., 2001; Pan et al., 1999; Yang et al., 2005).

In HD, extensive neuronal degeneration in major brain areas induce coordinated set of behavioural abnormalities in human patients as well as flies. Systemic metabolic derangements inflict additional burden on the health of already challenged individuals and gradually disables them. Although various strategies have been devised to slow down the disease progression, there are no effective treatment till now. Interestingly, curcumin with multiple cellular targets, minimum side effects and broad range of pharmacological activities can offer several advantages as a therapeutic choice over synthetic drugs with comparatively high toxicity and major side-effects.

Previously, the metabolic abnormalities seen in HD has been reiterated in *in vivo* transgenic *Drosophila* model expressing mutant human Htt (Aditi et al., 2016). In the present study, an effective concentration of 10μM curcumin administered to diseased flies since larval stages is shown to attenuate their motor dysfunction (Marsh et al., 2000; Marsh & Thompson, 2006), survival and metabolic abnormalities. We reinstated the effect of curcumin on locomotor functions in HD flies at an advanced stage (day 9) not reported earlier (Anjalika & Agrawal, 2016) (data not shown). This could be explained by the protection of motor neurons from mutant Htt-induced damage owing to the strong anti-oxidant property of curcumin. Curcumin feeding also improved the median survival in diseased flies which is again an indicator of enhanced protection from characteristic mutant Htt neurotoxicity and inflammation. Besides, curcumin-fed HD flies show fine management in abnormal fresh body weight as well as dry weight throughout the onset and progression of disease. However, once the disease becomes chronic, curcumin cannot mitigate them. It is possible that curcumin can recuperate the extent of neurodegenerative damages only upto a certain threshold of the disease. Curcumin targets or receptors might undergo progressive or deteriorative changes later and may become unavailable for further action (Phom et al., 2014).

Food intake quantification revealed that curcumin-fed HD larvae or flies did not display any prominent change in their feeding pattern which clearly indicated that weight modulation in these flies was not due to mere food intake but through modulation of metabolic process by curcumin. Amendment in dysregulated water, trehalose and lipid levels further corroborated the same. Water balance in *Drosophila* is usually regulated by neurosecretory or similar cell types, and any impairment in their functioning can lead to disproportionate water levels. Curcumin efficiently regulated the fluctuating water levels in HD flies and this might be attributed to better functioning of these cells in response to curcumin treatment. Additionally, curcumin effectively countered hyperglycemia in 7-day old HD flies. This effect could either be due to increased uptake of trehalose by cells or their controlled release into the circulation by source organs, apparently under the impact of curcumin. In contrast to these, no detectable effect of curcumin on other altered energy components *e.g.* protein or glycogen in HD flies was seen. The limited effect of curcumin on such critical energy reserves might explain the acute energy crisis prevalent at terminal stage when all the energy sources are urgently required for long-term sustenance and functional outcomes.

Curcumin exhibited prominent systemic and intracellular hypolipidemic effect in HD flies upon disease progression. The hypolipidemic effect might be due to the interaction of curcumin with important regulators of lipid metabolism. Several evidences point towards such beneficial effect of curcumin in attenuation of altered body weight, fat gain or glucose levels by regulation of multiple lipid metabolic genes (Ding et al., 2016; Shao et al., 2012). Based on these reports, we investigated the effect of curcumin on three key lipid regulators dSREBP, *bmm* and *lipin* in diseased flies, however surprisingly, no effect of curcumin on the expression of these genes in HD flies was seen. These findings indicate towards other critical players in lipid metabolism such as protein levels, activities, post-translational modifications, downstream effectors, target genes *etc*. a study of which will help elaborate the molecular action of curcumin in HD.

Chronic inflammatory conditions are often associated with neurodegenerative disorders and they can trigger numerous metabolic defects in the affected individuals. To elucidate the mode of action of curcumin in HD, its anti-inflammatory effect in HD flies was investigated. We noted an aggravated inflammatory state in HD flies, as denoted by the elevated ROS levels in adipose tissue, and 10μM curcumin effectively amended the abnormally high superoxide levels at both initial (day 7) and advanced (day 13) disease stages. Hence, the potent free-radical scavenging and anti-inflammatory property of curcumin proved immensely beneficial in attenuation of inflammatory and oxidative insult in HD thereby ameliorating the disease state.

## 5 CONCLUSION

Our findings show that curcumin is beneficial in suppression of neurodegeneration with amelioration of metabolic dysregulation. Though curcumin may not completely prevent neurodegeneration or metabolic impairments during terminal stages, it can effectively delay the inception and progression of HD at initial and moderate disease forms. Therefore, curcumin may prove to be a safe and suitable treatment regimen for management of HD that could be of great relief for the patients.

## AUTHORS’ CONTRIBUTIONS

KA, AS, MNS and NA perceived the idea, designed the research, interpreted and analysed the results. KA and AS performed the experiments and carried out data collection. KA and AS wrote the manuscript. All authors read, reviewed and approved the final manuscript.

## ACKNOWLEDGEMENTS

We thank Prof. J. Lawrence Marsh, University of California Irvine, CA, USA for providing transgenic *Drosophila* lines *UAS-Httex1p Q20*, *UAS-Httex1p Q93* and *elav-GAL4* driver. We are thankful to Dr. Ekta Kohli, Defense Institute of Physiology and Allied Sciences (DIPAS), Defense Research and Development Organisation (DRDO), Delhi, India for providing RT-PCR facility. KA and AS acknowledge University Grants Commission (UGC) and Council of Scientific & Industrial Research (CSIR), respectively, for providing fellowship to conduct research.

